# Physical–Chemical Approach to Identify Local Structural Determinants of Molecular Mechanisms: Case Study of Antimalarial Drug Pyronaridine and Crystal-Growth Inhibition

**DOI:** 10.1101/2025.11.04.686507

**Authors:** Angela Medvedeva, Ksenia Kolomeisky, Anatoly B. Kolomeisky

## Abstract

Understanding how specific molecular substructures control chemical behavior is central to rational molecular design and the development of new materials. However, most current predictive models offer limited mechanistic resolution at the fragmental level. We present a novel physical–chemical computational framework, which is based on systematically perturbing a parent molecule, to quantify fragment-level contributions to both a specific mechanistic action and a broader functional outcome. To test our theoretical approach, we investigated local structural contributions to pyronaridine (PY), a clinically used antimalarial drug with a mechanistically distinctive mode of inhibition of hematin crystal growth via step-bunching. Chemically plausible PY molecular analogs have been computationally generated by selectively removing or substituting functional groups hypothesized to influence either step-bunching mechanisms or whole-parasite blood-stage activity. For each analog, we predicted the probability of four different crystal-growth inhibition mechanisms using a centroid-based similarity model based on a small dataset of experimentally verified crystal-growth inhibitors. The blood-stage antimalarial activity has also been estimated using the MAIP platform. A systematic comparison of molecular analogs revealed that step-bunching mechanisms depend primarily on two protonated pyrrolidines, with chlorobenzene as a strong secondary contributor. In contrast, antimalarial activity is more distributed, relying on coordinated interactions between aromatic–heteroatom scaffolds and an amine linker. The obtained results demonstrate that our approach can disentangle position-specific and cooperative fragmental effects, offering mechanistically interpretable guidance for the design of mechanism-optimized inhibitors. The framework might be broadly applicable across chemical and materials domains where linking local structure to specific mechanisms is essential.

## I. INTRODUCTION

The development of novel medicines and materials critically depends on the ability to connect the overall functionality with the properties of specific molecular fragments.^1^ Yet, current methods in materials chemistry and drug discovery, such as machine-learning quantitative structure–activity relationship (QSAR) models that map molecular structure to activity, can often predict whether a molecule is likely to be active, but not which specific substructures matter most.^2^ Experimental structure–activity studies have made important progress by investigating amino acid substitutions in peptides^3^ and by modifying small molecules one fragment at a time, including crystal-growth modifiers used to treat cystinuria^4^ and malaria.^5^ However, even small changes can produce unexpected behavioral shifts,^5^ and experimentally mapping these relationships to molecular structures can be difficult and resource-intensive.

Decades of quantitative structure–activity relationship (QSAR) research have delivered robust predictors across medicinal and materials chemistry.^2,6^ Frequently, the focus is on global molecular descriptors or fragment-encoded fingerprints that correlate the overall molecular features with outcomes, powerful for prediction but often still limited for substructure-level mechanistic understanding. Classic methods such as Free–Wilson analysis^7,8^ and fragment-based QSAR approaches, including hologram QSAR models,^9–13^ estimate additive fragment contributions across series of chemical molecules,^14,15^ while matched molecular pair analysis (MMPA) studies dataset-wide chemical substitutions to understand property shifts.^16–18^ Molecular fragmentation strategies are also widely applied for similarity searching^19^ and for decomposing molecules in generative or property-prediction models^20^. More recently, explainable-AI toolkits such as SHAP^21,22^ and graph neural network (GNN) substructure-masking^23,24^ have provided local attributions for complex models. These approaches are valuable for identifying statistical associations, highlighting informative correlations, and generating counterfactual examples.^25^ Nonetheless, they mostly clarify global correlations: while they can suggest which features are associated with a specific outcome, they rarely directly test the specific contribution of predefined substructures to a specific predicted mechanism or behavior.

It is widely accepted that global molecular properties, such as crystallization outcomes, mode of transport, or bioactivity, are ultimately governed by local structural features and their interactions. We hypothesize that (i) discrete substructures make mechanism-specific contributions, and (ii) combinations of substructures can display non-additive (synergistic or redundant) effects that shape distinct global behaviors. Even if these local effects can be estimated only indirectly, such measurements can offer an interpretable route to mechanistic insights and rational design, particularly for small-data or rare-mechanism regimes, where conventional QSAR approaches may struggle due to limited training coverage.

To test our hypothesis, we introduce a minimal interpretable strategy to identify substructures in a given molecule that most strongly associate with a target behavior, provided that the behavior can be predicted by any reasonable model. This approach provides a framework for understanding, in principle, the behavior of any molecule as a combination of molecular fragments. Starting from a parent molecule, we systematically remove functional groups or fragments *in silico* to create a panel of chemically reasonable analogs, predict each analog’s behavior, compare the predictions to the parent molecule, and attribute the largest changes to the removed substructures. Because the method operates directly on chemically meaningful substructures rather than abstract descriptors, it yields fragment-level mechanistic insight at the level that is typically utilized in chemistry. The framework is also predictor-agnostic: any computational outcome (physical, chemical, or biological) can be easily utilized in the analysis.

To illustrate our approach, we apply it to a clinically relevant and mechanistically well-quantified case of the pyronaridine (PY) molecule, which is an antimalarial drug associated with the so-called step-bunching mode of hematin crystal-growth inhibition. Using a custom model trained on a small but well-characterized dataset, the probability of one of four currently known crystal-growth inhibition mechanisms, including also a non-inhibitory category, is estimated. We connect it with the predictions from the MAIP antimalarial activity tool^26^ to compare fragment contributions to overall blood-stage antimalarial activity versus those specific to the step-bunching mechanism. This dual-predictor design enables us to separate mechanism-critical structural features from those that contribute more broadly to whole-parasite neutralization.

Malaria provides a biologically and clinically important test case where such structure–mechanism mapping is directly relevant. In the blood stage of infection (outlined in detail in the Supporting Information), *Plasmodium* parasites digest host hemoglobin within an acidic digestive vacuole, releasing ferriprotoporphyrin IX (Fe(III)-heme), a redox-active and membrane-disruptive toxin^27,28^. To survive, the parasite crystallizes heme into insoluble hemozoin crystals^29,30^. Many antimalarial drugs act, at least in part, by disrupting hemozoin crystal growth, thereby increasing toxic free-heme levels and killing the parasite.^5,31–34^ Crystal growth proceeds through the lateral motion of molecular “steps” across the crystal faces; if these steps accumulate unevenly, which is known as step-bunching, the surface jams, and growth slows down or even stops. Although step-bunching is a well-established mechanism for crystals, it is still a less common scenario compared to other crystal-growth phenomena.^35^ Among small-molecule drugs, it remains under-characterized in terms of structural determinants at biological interfaces.

Pyronaridine (PY) is a clinically used antimalarial medicine, which is one of the few small molecules experimentally associated with step-bunching in hematin crystallization with high confidence^36^. This makes PY an ideal proof-of-concept system for our framework: it is mechanistically distinctive, biologically relevant, relatively simple chemically, and it exhibits a growth-inhibition mode that is both rare and interpretable. By pairing a custom step-bunching probability model with the established MAIP blood-stage antimalarial activity predictor, we use PY to (i) identify fragments with primary, position-specific roles in step-bunching and/or antimalarial activity, (ii) determine which fragments are selectively important for one endpoint versus the other, and (iii) map additive and non-additive fragment interactions.

## II. METHODS AND MATERIALS

### A. General Approach

A general overview of our computational approach is presented in Fig. 1. Pyronaridine (PY) is used as the parent compound for the analysis. Two predictions of chemical behavior are generated for PY and for a series of systematically modified analogs: (1) probability of step-bunching in hematin crystal growth, and (2) predicted antimalarial activity using the MAIP platform.^26^

**FIG. 1:**
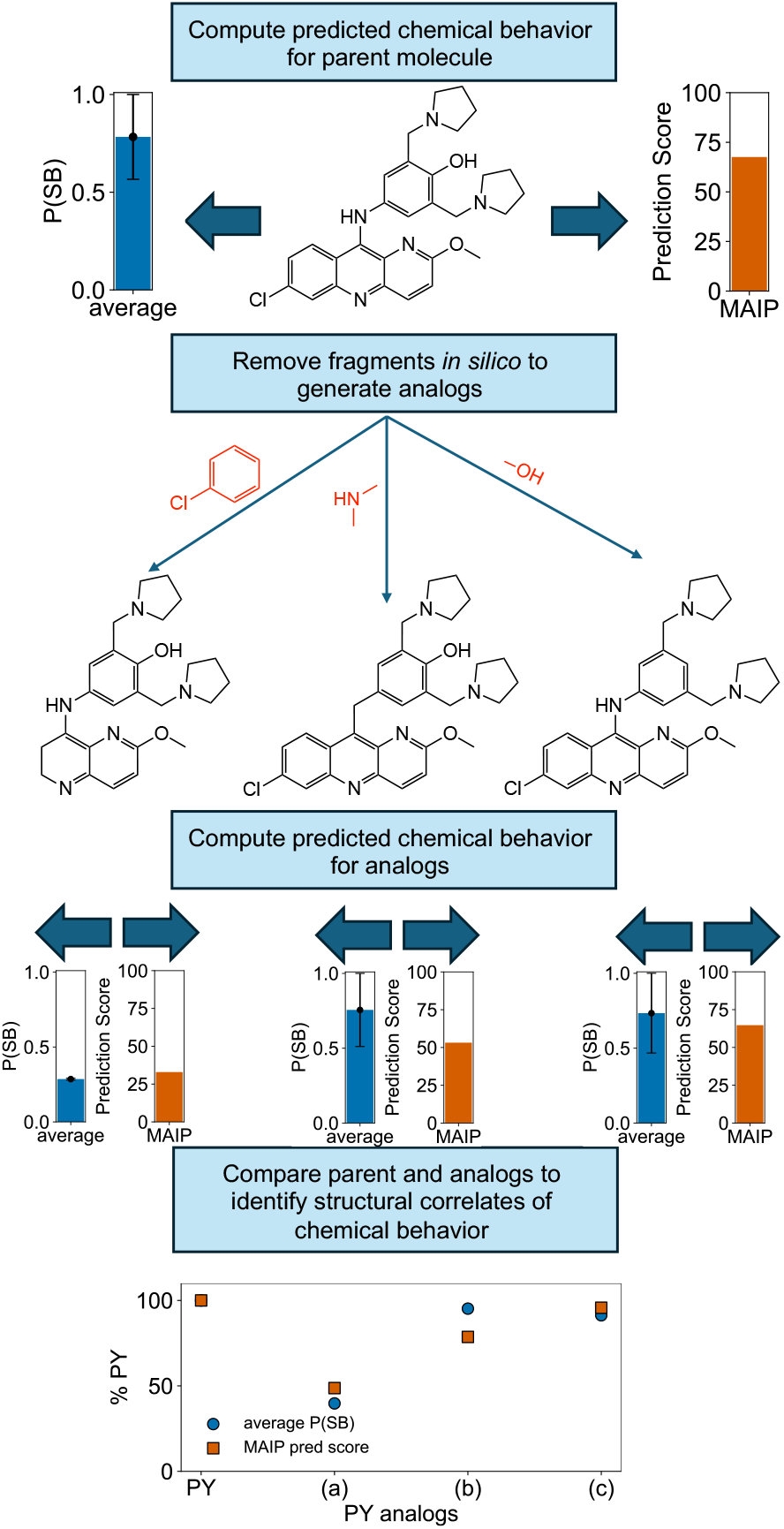
Computational approach overview. The parent compound (PY) is analyzed to estimate the mean probability of step-bunching mechanisms (P(SB)) and MAIP-predicted antimalarial activity (MAIP). Structural fragments are then selectively removed or altered to generate analogs. Analog predictions are compared to the parent molecule to assess the contribution of individual fragments to each behavior.

### B. Crystal-Growth Inhibition Mechanism

Crystals grow by adding atomic or molecular steps across their surfaces.^37,38^ Small molecules can modulate the growth dynamics through several distinct, surface-specific inhibition modes, each defined by the geometry and persistence of its effect on the step propagation.^36^ Accordingly, such molecules are known as crystal-growth *modifiers* or inhibitors.

There are four possible mechanisms of how small molecules might influence crystal growth. The first is *step-pinning (terrace/step adsorption)*.^36,39,40^ In this case, the modifier molecules adsorb at terraces or along step edges, creating fixed “pinning” points that force growing steps to bow around the obstacles. This slows down layer-by-layer growth, but it typically remains reversible once the modifier is removed. Many quinoline antimalarial drugs exhibit such step-pinning behavior in blocking hematin crystallization.^41^

Another inhibition scenario is known as *kink-blocking (site poisoning)*.^5,36,42^ Here, modifiers bind directly at kink sites—the high-energy points where growth units preferably attach during step advance—preventing incorporation of new molecular units into crystal until the inhibitor desorbs or new kinks form. This site-specific poisoning produces a more complete local arrest than the step-pinning mechanism, but is spatially restricted to kink regions, and it is also generally reversible.

A more complex mechanism is *step-bunching (cooperative step instability)*.^36,43–46^ In this case, rather than slowing isolated steps, modifiers perturb the step train as a whole, causing adjacent steps to coalesce into “bunches.” These bunches can seed dislocations and other long-lived defects, permanently reducing growth rates even after inhibitor removal^43^. Step-bunching is therefore a cooperative, history-dependent mechanism with irreversible consequences for surface morphology. Pyronaridine in hematin crystallization^36,43,44^ and certain L-cystine mimics in blocking cystine crystals^45,46^ are well-characterized examples of such a crystal-growth inhibition mechanism.

The final possibility is known as the *passive-bystander* mode.^43,47^ In this scenario, the compound associates with the crystal surface without meaningfully altering step advancement rates or morphology under experimental conditions. Binding may be transient, weak, or incorrectly oriented for interference with key growth sites. Crystal-growth inhibition does not occur in this mode of action. In this study, we evaluate the probability of each of these four mechanisms—step-pinning, kink-blocking, step-bunching, and no crystal-growth inhibition (passive-bystander)—for pyronaridine (PY) and for each designed analog. Probabilities are estimated using a custom centroid-based scoring method (detailed in the Supporting Information) that compares each molecule to a small, experimentally verified reference set of mechanism-annotated hematin growth inhibitors.

For our specific calculations, the mean probability of the step-bunching mechanism (P(SB)) is used as the primary quantity of interest for the crystal growth mechanism outcome. It is computed using two complementary approaches, *inverse-distance normalization*, which yields higher probabilities for the closest class, aiding interpretability; and *softmax transformation*, which produces more balanced distributions, reducing the influence of small absolute distance differences. The reported mean P(SB) is the average of the two methods, with the standard error (SE) given as half the range between them. Because the model operates in a similarity-based, clustering framework rather than as a fully trained classifier, the absolute values should be interpreted qualitatively, with emphasis on relative changes across analog series rather than definitive class assignments.

The predicted antimalarial activity is computed using the MAIP (MAlaria Inhibitor Prediction) web tool,^48^ an open-access, consensus machine-learning platform hosted by EMBL–EBI that integrates QSAR models from multiple proprietary and public datasets to identify small molecules with blood-stage *Plasmodium falciparum* (one of the parasites that causes malaria) inhibition potential. The current model is specifically trained to predict activity against this *Plasmodium falciparum* parasite and outputs a continuous regression-like score representing the predicted blood-stage antimalarial activity. Higher scores indicate a greater likelihood of blood-stage activity, but the score does not represent a probability. Because the output is unbounded and non-probabilistic, scores are relative rather than absolute. Consequently, the score for an untested analog should be interpreted in relation to that of a known antimalarial with established blood-stage antimalarial activity, such as PY.^26,48^ MAIP has been experimentally validated to increase hit rates by up to 12-fold compared with the random screening in *Plasmodium falciparum* blood-stage assays.^26^ Discussion on the significance of the targeted *Plasmodium* species and life-cycle stage of inhibition is provided in the Supporting Information.

### C. Parent Molecule: PY as a Model System

Pyronaridine (PY) is a convenient model system for our investigation for several reasons. First, understanding how PY induces step-bunching is relevant for clarifying the fundamental aspects of crystal growth and inhibition.^49,50^ Small-molecule crystal-growth inhibitors are intrinsically valuable in this context because they can be engineered to bind specific targets, enabling precise mechanistic tests and control of mechanisms. In medicine, such molecules are rare, but they are quite impactful, with notable examples in malaria (heme crystallization inhibitors)^43,44^ and cystinuria (L-cystine mimics that alter cystine stone growth).^45,46^

Second, PY’s inhibition of hematin crystallization is also an exceptional microscopic phenomenon. The molecule not only binds to crystal surfaces but also induces step-bunching – a cooperative, history-dependent, and irreversible process that produces persistent lattice defects, halting growth even after drug removal.^43,44^ This sets it apart from the other two main inhibition modes: step pinning, where adsorbed molecules slow step advance in a reversible manner, and kink blocking, where growth units are prevented from attaching at active sites until the inhibitor is removed.^36^ The permanence of PY-induced lattice disruption, combined with potent nucleation suppression, provides a rare opportunity to study how multi-modal molecular interactions (*π*–*π* stacking, hydrogen bonding, metal coordination) might collectively control crystal morphology and stability at the nanoscale.^44,51,52^

Third, there are direct translational applications of understanding PY’s local structural features for drug development. Artemisinin-based combination therapies (ACTs) are the current frontline treatment for malaria, pairing a fast-acting but short-lived artemisinin derivative with a slower-eliminating partner drug to ensure clearance of residual parasites and delay resistance.^53^ Artemisinin, once a dominant monotherapy before the emergence of resistance, is not a crystal growth inhibitor; it most probably acts through endoperoxide activation by heme iron to kill parasites rapidly, and is now used only in ACTs.^54^ PY’s ability to act at multiple lifecycle stages, including ring-stage artemisinin-resistant parasites,^44^ makes it particularly effective as an ACT partner^55,56^. Sustaining the clinical efficacy of combination therapies requires understanding PY’s mechanistic compatibility with other inhibitors, which in turn depends on elucidating the structural determinants underlying its antimalarial action.

Fourth, studies of binary combinations of crystal growth modifiers show that pairs with distinct mechanisms (e.g., step pinners like chloroquine and kink blockers like amodiaquine) can produce not only synergy but also antagonistic cooperativity, where one inhibitor reduces the effectiveness of the other by altering surface properties.^40^ PY’s unique step-bunching mechanism provides a mode of hematin inhibition distinct from the step-pinning and kink-blocking profiles of most other quinoline-class ACT partners, which may influence combination performance and offer a scaffold for exploring mechanistic complementarity in ACT partner selection. PY is also one of the very few small-molecule inhibitors in any disease system (alongside with certain L-cystine mimics in cystinuria^57^) for which high-resolution atomic force microscopy (AFM) studies have established an unequivocal mode of action as a step-buncher.^36,46^ This rare combination of fundamental scientific relevance, proven clinical utility, and mechanistic distinctiveness makes PY a special model for bridging crystal growth science with urgently-needed therapeutic innovation.

### D. Analog Design Rationale

In our prediction of crystal growth inhibition mode, Pyronaridine (PY) is contrasted with canonical quinoline antimalarial crystal-growth inhibitors such as chloroquine (CQ) and quinine (QN) that follow the step-pinning mechanisms, and with amodiaquine (AQ) and mefloquine (MQ) that follow the kink-blocking mechanism.^36^ Fig. 2 presents the molecular structures of PY and other antimalarial drugs in our dataset (detailed in the Supporting Information), highlighting the key functional groups present in each molecule and summarizing their proposed roles based on previous studies. Compared to chloroquine (CQ), quinine (QN), amodiaquine (AQ), and mefloquine (MQ), PY uniquely incorporates nearly the full set of potentially functional fragments observed across these crystal-growth inhibitors, including aromatic rings, protonatable amines (secondary and tertiary), halogens, ethers, hydroxyls, and phenols.^56^ In contrast, CQ lacks an ether and a hydroxyl, QN lacks halogens, MQ lacks ethers, and CQ, QN, and MQ all lack a phenol group.

**FIG. 2:**
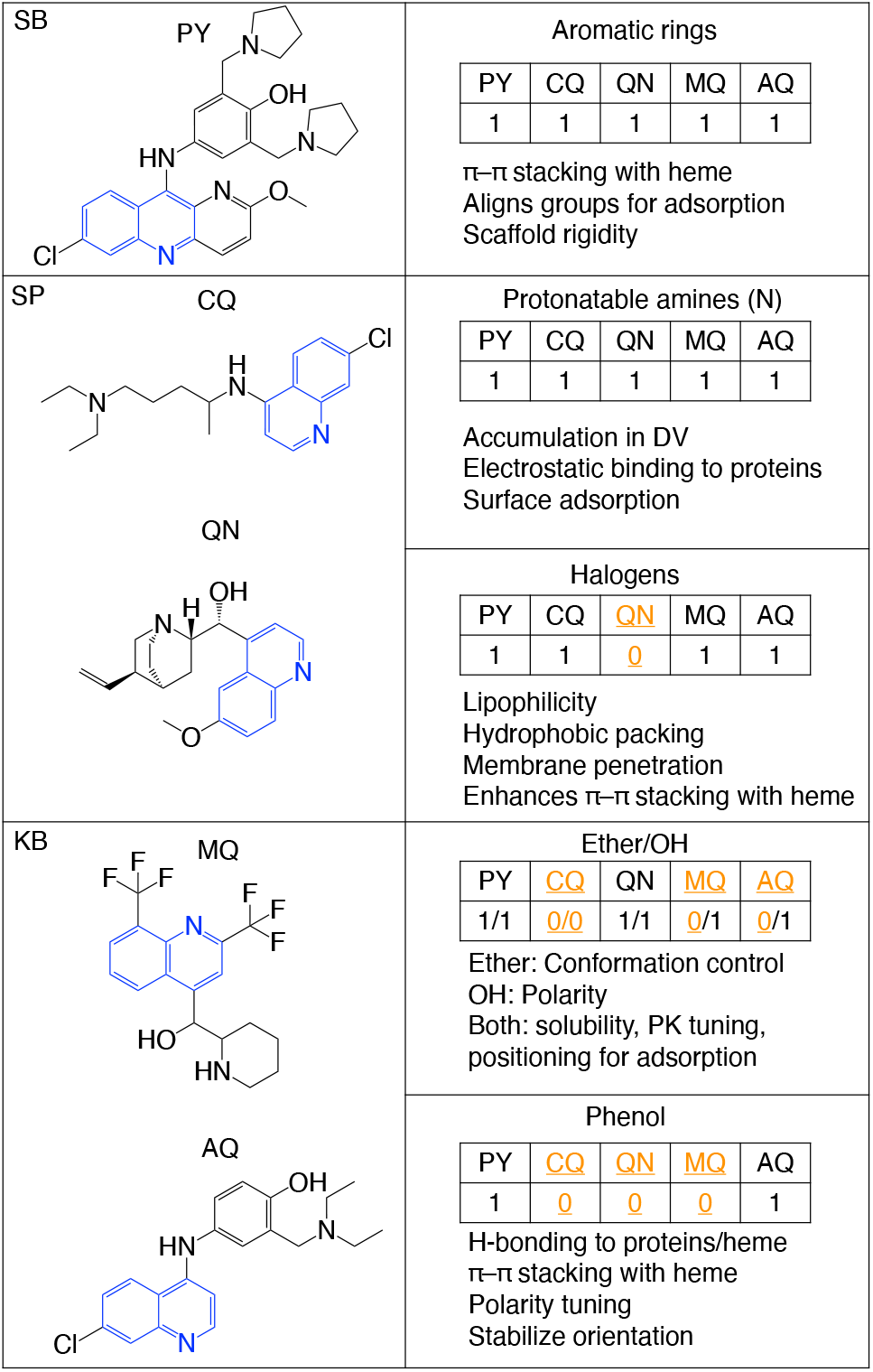
Molecular structures of step-buncher (SB) pyronaridine (PY), step-pinners (SP) chloroquine (CQ) and quinine (QN), and kink-blockers (KB) mefloquine (MQ) and amodiaquine (AQ), with key functional groups present in each compound indicated in the accompanying tables (presence = 1; absence = 0, shown in orange and underlined), and proposed roles of those groups based on prior studies. The quinoline core, shared by all crystal growth inhibitors in the dataset, is shown in blue in each molecular structure.

All crystal-growth inhibitors in the dataset share a quinoline core (shown in blue in Fig. 2). It is believed that aromatic rings, such as those in the quinoline core, contribute to *π*–*π* stacking with the porphyrin core of heme, which aligns hematin molecules for optimal adsorption to crystal faces, imparting also the scaffold rigidity to maintain interaction geometry.^39,58^ These properties arise from the planar, conjugated nature of aromatic systems, allowing them to interact closely with the flat porphyrin ring and maximizing van der Waals and *π*-electron overlaps. By acting as a rigid framework, aromatic rings also ensure that other functional groups, such as amines or hydroxyls, are presented in the orientations favorable for simultaneous binding events at the crystal interface.

The amine groups, which are often protonated in the acidic digestive vacuole of the malaria parasite, facilitate acid trapping, electrostatic binding to heme propionates, and adsorption to crystal surfaces.^39,41,60–62^ Protonation converts these amines into cationic centers, which are electrostatically attracted to the negatively-charged propionate groups on the heme. This effect not only anchors the compound to the crystal face but also increases local concentration inside the vacuole through pH-driven retention, amplifying the potential for inhibitory interactions.

Halogens can enhance lipophilicity (i.e., the affinity for a non-polar environment), promote hydrophobic packing and membrane penetration, and strengthen *π*–*π* stacking interactions with heme.^63,64^ These effects result from the halogen’s size, polarizability, and ability to withdraw or donate electron density, thereby modulating the *π*-system’s interaction strength with the aromatic surface of heme. Increased lipophilicity also aids in crossing lipid membranes to reach the digestive vacuole.

Ethers can lock parts of the molecule into conformations that favor surface binding,^65–67^ while hydroxyls form hydrogen bonds with surrounding solvent, protein residues, or heme propionates.^68–70^ Together, ether and hydroxyl groups can modulate solubility, kinetics of association, and positioning for effective adsorption.^71–73^

Phenolic *OH* (*Ar*–*OH*) groups differ from aliphatic *OH* (*R*–*OH*) in their greater acidity and resonance stabilization, enabling stronger and more directional hydrogen bonding to proteins or heme, *π*–*π* stacking enhancement, polarity tuning, and stabilization of molecular orientation at the heme surface.^74–76^ The conjugation between the phenolic oxygen and the aromatic ring stabilizes the phenoxide anion formed upon hydrogen bond donation, which can lead to stronger and more geometrically-precise interactions at the crystal interface. When positioned near an aromatic core, phenolic *OH* can also synergize with *π*–*π* stacking, locking the molecule into an orientation that maximizes inhibitory contact with hematin surfaces.

It has been reported^77^ that stronger crystal growth inhibition often results from a combination of *π*–*π* stacking capacity (extended aromatic systems) and direct *Fe*(III) coordination through donor atoms such as phenolic or heteroaromatic nitrogens. Within this dual-mode framework, PY’s phenol, hydroxyl, ether, and multiple amine functionalities may form a multifunctional scaffold capable of simultaneous hydrogen bonding, metal coordination, and aromatic stacking, with the phenol providing a directional *H*-bonding anchor. Certain groups (e.g., aromatic nitrogen donors or phenolic *OH*) are likely to act directly at crystal-growth steps via face binding or step bunching, while other groups tune geometry, solubility, and molecular orientation, thereby amplifying the effectiveness of the primary interactions.

Taking into account the above considerations about the properties of different fragments, we manually designed a set of 25 chemically reasonable pyronaridine (PY) analogs in ChemDraw (Revvity Signals Software) to systematically probe fragment-level contributions. The analog library was constructed to remove or substitute specific functional groups, including amines/tertiary nitrogens, the halogen, ether, and hydroxyl groups (with explicit contrasts between phenolic and aliphatic *OH*). In addition, our goal was to perturb the PY scaffold to test whether its saturated ring content and functional-group multiplicity are necessary for the step-bunching mechanism of crystal growth inhibition versus broader (whole-parasite) antimalarial activity. This design allowed targeted evaluation of the relationship between defined structural elements and two distinct outcomes: (1) step-bunching probability and (2) blood-stage antimalarial activity. All structures were exported as SMILES text strings and submitted to both the step-bunching probability prediction model and the MAIP antimalarial activity prediction tool.

For each pyronaridine analog, the predicted step-bunching mechanism probability was computed using the step-bunching probability prediction method. In parallel, predicted blood-stage antimalarial activity was obtained for each analog using the MAIP predictor. The analogs were then compared to the parent in each metric, allowing a direct, side-by-side comparison of fragment-level effects on a specific crystal-growth inhibition mechanism versus whole-parasite activity. These comparisons formed the basis for classifying each fragment as having a primary, secondary, or supporting role in step bunching and/or broader antimalarial activity.

Analog performance was compared to the parent PY in two ways: (1) as a percentage of the parent value (%PY), and (2) as percent reduction relative to the parent molecule (%Δ).

## III. RESULTS AND DISCUSSION

We start with the chemical structure and properties calculated using our approach for the parent molecule, pyronaridine (PY), as presented in Fig. 3. The structure of PY is shown in Fig. 3(a). One can estimate the mean probability of inhibition of crystal growth via the step-bunching mechanisms, and it is predicted to be P(SB)=0.78, as shown in Fig. 3(b) on a scale from 0–1 scale. In parallel, the MAIP prediction tool estimates PY’s blood-stage antimalarial activity at MAIP = 67.57. Although this value is plotted on a 0–100 scale in Fig. 3(c) for ease of visualization, the MAIP score is a relative, unbounded quantity rather than a true probability. PY has been experimentally confirmed to exhibit both step-bunching behavior and blood-stage antimalarial activity. These predicted values therefore represent an approximate quantitative description associated with the emergence of these behaviors. Our idea is that analogs with values that deviate substantially from these thresholds are less likely to display comparable step-bunching or antimalarial activity. Therefore, by examining the fragments that distinguish PY from its analogs, we can infer which structural features are associated specifically with step-bunching, antimalarial activity, or both, based on reductions in the corresponding predicted values for the analogs.

**FIG. 3:**
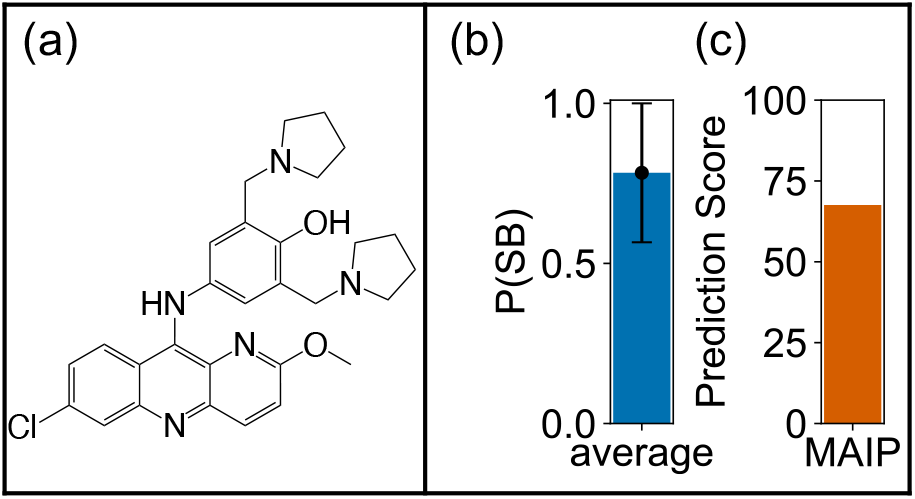
(a) Chemical structure of pyronaridine (PY); (b) mean predicted probability of step bunching, with error bar showing half the range from two different prediction methods; and (c) predicted blood-stage antimalarial activity from the MAIP prediction tool.

Fig. 4 shows the effect in predicted step-bunching probability and antimalarial activity, relative to the original PY molecule, for a series of analogs generated by sequential fragment substitution or removal. The leftmost vertical column shows the fragments that were removed or substituted in red, and the center vertical column shows the resulting analogs. The sequence of analogs begins with the substitution of nitrogens (*N*) in the pyrrolidine rings of PY with *CH* groups (panel a), followed by the removal of both 5-membered rings and their ethyl linkers from PY (panel b), the panel (b) analog with the additional removal of the phenol group (panel c), and finally the panel (c) analog with additional removal of the amine linker (panel d). Panel (e) summarizes each analog’s predicted step-bunching probability and antimalarial activity as a percentage of the original PY values.

**FIG. 4:**
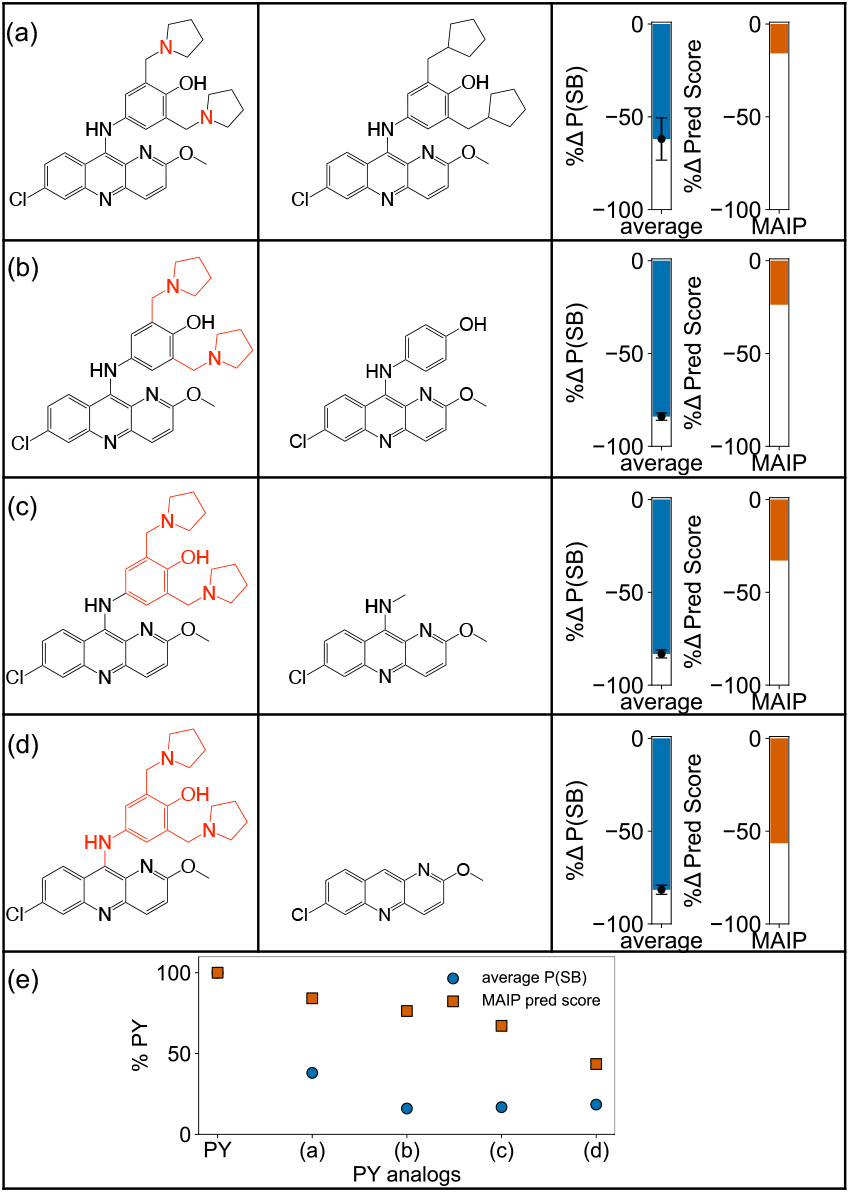
Predicted impact of sequential fragment substitution or removal from pyronaridine (PY) on probability of step-bunching mechanisms and antimalarial activity. (a) Substitution of pyrrolidine nitrogens (*N*)with *CH* groups; (b) removal of both pyrrolidines and their ethyl linkers; (c) the same as in (b) but with additional removal of the phenol group; (d) the same as (c) but with additional removal of the amine linker. (e) Predicted values expressed as a percentage of PY values. The left vertical column shows in red the chemical entities removed for creating PY analogs. The middle vertical column shows the considered analogs. Error bars represent the range-based variation between two independent step-bunching probability prediction methods.

Our computational analysis (see Fig. 4) suggests that the largest reduction in step-bunching probability occurs upon the removal of both 5-membered rings and their ethyl linkers (*−*84.0%, panel b). Similar reductions have been observed for the removal of the phenol group (*−*83.2%, panel c) and the amine linker (*−*81.6%, panel d). Substituting only the pyrrolidine nitrogens (*N*) with *CH* (panel a) resulted in a smaller, though still substantial reduction (*−*62.0%). The smaller effect in (a) underscores the importance of the pyrrolidine nitrogens (*N*) within the 5-membered rings; systematic nitrogen substitutions (see Fig. S1) confirm that these nitrogens, particularly in the pyrrolidine rings, are probably most strongly associated with the step bunching mechanism. Removing even a single pyrrolidine with its ethyl linker, as shown in Fig. S1(b), causes a substantial reduction (*−*69.2%), though this is still less than the reduction from removing both (*−*84.0%), shown in Fig. 4(b). As shown in Fig. 4(e), additional removal of structures near the 5-membered rings (as in analogs c and d) did not further strongly reduce predicted step-bunching, suggesting that these fragments probably serve mainly as scaffolds.

In terms of predicted antimalarial activity, the MAIP score decreased most with the largest extensive fragment removal (see Fig. 4, panel d). As shown in panel (e), the decline was approximately linear with increasing fraction of fragment removals, indicating that each fragment contributes uniquely to the predicted antimalarial activity. However, this trend should be interpreted with caution, as the decreases might reflect the cumulative loss of molecular components rather than the specific removal of the most functionally important fragments. To identify the role of nitrogens in other aspects of antimalarial activity, Fig. S1 further analyzes nitrogen atom substitutions one at a time. This approach minimizes confounding effects related to fragment size or overall molecular mass, thereby clarifying the contributions of specific functional groups. It shows that the predicted antimalarial activity was lowest for the amine linker, with no reduction in any of the other analogs.

Fig. 5 shows the effect of different approaches to creating analogs. Removal of the chlorobenzene group from PY resulted in a 60.3% reduction in the predicted step-bunching probability and a 51.3% reduction in the predicted antimalarial activity (panel a). Subsequent removal of the pyridine and the ether group from analog (a) led to slightly greater reductions in both predicted step-bunching probability and predicted antimalarial activity (*−*63.4% and *−*59.4%, respectively) (panel b). Further removal of the pyridine linked to the amine linker from analog (b) produced a similar step-bunching reduction (*−*63.9%) and antimalarial activity (*−*58.7%) (panel c). The most extensive fragment removal, which eliminated the amine linker from analog (c), yielded, as expected, the largest reductions: –69.2% for step-bunching probability and –69.8% for antimalarial activity (panel d).

**FIG. 5:**
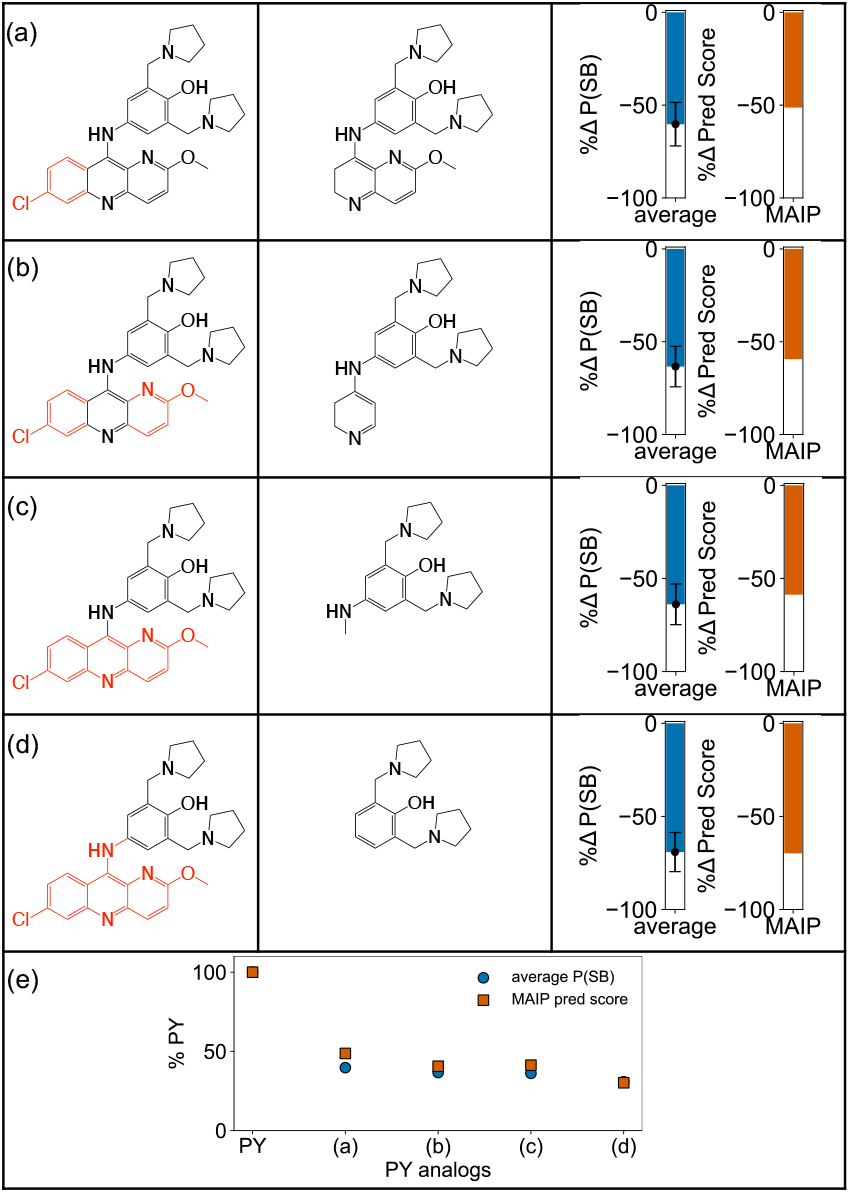
Predicted step-bunching probability and antimalarial activity for pyronaridine (PY) analogs with sequential fragment removal. (a) Chlorobenzene removed; (b) the same as in (a) with additional removal of the pyridine and ether; (c) the same as in (b) with additional removal of the pyridine linked to the amine; (d) the same as in (c) with additional removal of the amine linker. (e) Predicted values expressed as a percentage of PY metrics. The left vertical column shows in red the chemical entities removed for creating PY analogs. The middle vertical column shows the considered analogs. Error bars represent range-based variation between two independent step-bunching probability prediction methods.

Panel (e) expresses each analog’s predicted values as a percentage of the original PY properties. Across all the analogs, further fragment removals after chlorobenzene elimination produced less than a 10% additional reduction in the step-bunching probability and less than a 20% additional reduction in the predicted antimalarial activity. This result indicates that chlorobenzene is probably the dominant single contributor among the considered fragments, and it also suggests that it may exert an independent effect beyond synergistic interactions with other structural elements.

Chlorobenzene and pyrrolidines were independently identified as key fragments in separate analyses, based on the observation that further removal of adjacent functional groups or heteroatoms after eliminating either fragment had little additional impact on the reduced step-bunching probability or antimalarial activity. These observations suggest that the remaining molecular components serve primarily as a supporting scaffold. Our analysis indicates that both the chlorobenzene and the two pyrrolidines with their ethyl linkers are strongly associated with step bunching (see also Fig. S2), with chlorobenzene also moderately linked to antimalarial activity. Overall, these results suggest non-additive effects from different fragments for step-bunching, in which the primary contribution comes from the pyrrolidines. Fig. S2 suggests possible synergistic effects for antimalarial activity, where both fragment types contribute substantially when combined or removed together.

Fig. 6 shows the results of our analysis for individual functional group modifications in PY. Removal of only the hydroxyl group (panel a), removal of only the ether (panel c), and removal of only the chlorine atom (panel d) from PY each resulted in predicted values remaining above 89.0% of those for PY, for both step bunching and antimalarial activity. Substitution of the amine in the linker (*NH*) with a *CH*_2_ group (panel b) also had little effect on the predicted step-bunching probability but produced a notable 21.4% decrease in predicted antimalarial activity, to 78.6% of PY’s value. This suggests that the linker amine is more strongly associated with antimalarial activity than other individual functional groups examined.

**FIG. 6:**
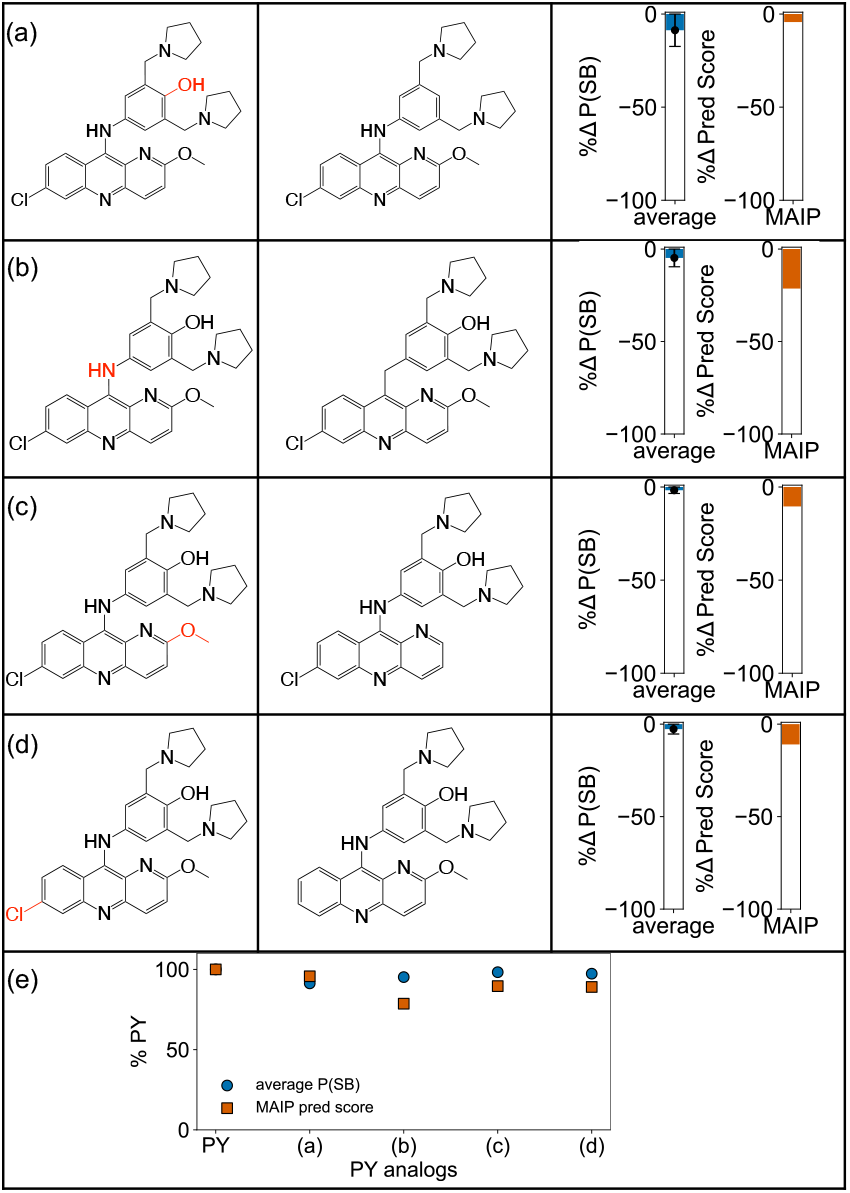
Predicted step-bunching probability and antimalarial activity for pyronaridine (PY) analogs with individual functional group modifications. (a) Hydroxyl group removed; (b) amine in linker (*NH*) substituted with *CH*_2_; (c) the ether group removed; (d) chlorine removed; (e) predicted values expressed as a percentage of PY. The left vertical column shows in red the chemical entities removed for creating PY analogs. The middle vertical column shows the considered analogs. Error bars represent range-based variation across two different step-bunching probability prediction methods.

Fig. 7 summarizes the overall analysis of the fragment–function relationships, combining fragments according to the hypothesized functional categories in Fig. 2 and classifying each as having a primary, secondary, supporting, or minor role in step-bunching and/or antimalarial activity. Below we discuss in detail the proposed roles of different fragments and groups in the supporting step-bunching mechanisms and the overall antimalarial activities.

**FIG. 7:**
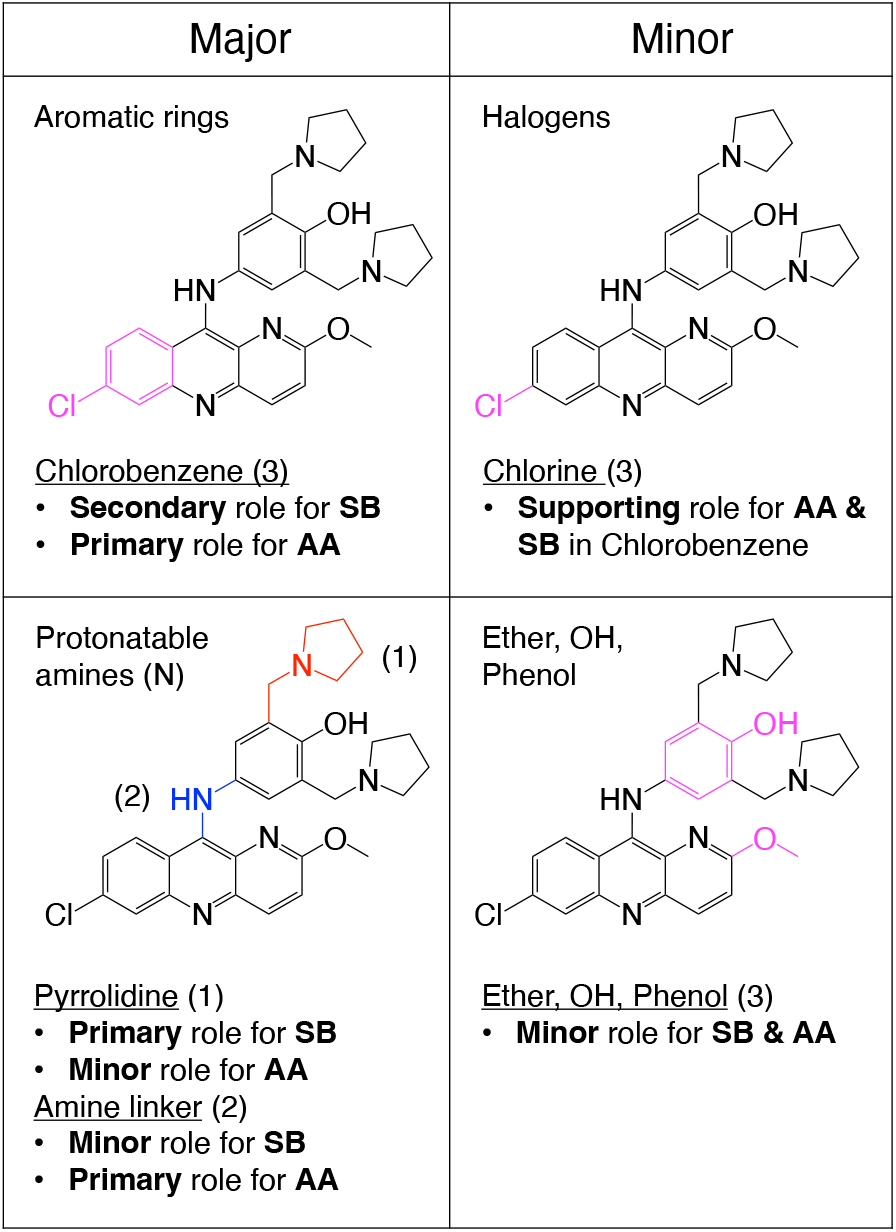
Summary of functional group categories and specific fragments tested in step-bunching (SB) and antimalarial activity (AA). Fragments are grouped by their broader hypothesized functional categories (Fig. 2) and classified according to the magnitude and specificity of their effect on predicted SB and AA values. “Major” fragments produce large, independent effects on one or both mechanisms, whereas “minor” fragments have smaller or supporting effects, typically acting in combination with other groups. Color and numerical coding correspond to the text: magenta and (3) denote fragments with similar significance for SB and AA, red and (1) denote SB-specific significance, and blue and (2) denote AA-specific significance.

### A. Major roles: aromatic rings and protonatable amines

It has been shown before that aromatic rings contribute to *π*–*π* stacking with the porphyrin core of heme, align groups for optimal adsorption to hematin crystal faces, and increase the scaffold rigidity to maintain interaction geometry.^39,58,59,78,79^ Amines, often protonated in the acidic digestive vacuole, facilitate acid trapping, electrostatic binding to heme propionates, and adsorption to crystal surfaces.^39,41,60–62^

Our analysis suggests that among all aromatic features in PY, the chlorobenzene ring emerges as the key contributor. Removal of chlorobenzene significantly reduced step-bunching probability and antimalarial activity (Fig. 5a), making it the largest single-fragment determinant of antimalarial activity and a strong secondary contributor to step-bunching. In contrast, when chlorobenzene and pyrrolidines are left intact but all other aromatic rings are converted to saturated analogs (see Fig. S3b), step-bunching is essentially unchanged, while antimalarial activity falls by 30.8%. This indicates that for step-bunching, peripheral aromatic scaffolds play a minor role as long as the primary fragments remain, whereas antimalarial activity depends more on the integrated contributions of multiple aromatic regions.

For step-bunching, the actual *position* of the aromatic group matters. Introducing aromaticity into the pyrrolidines by converting them to pyridines (see Fig. S3c) does not significantly reduce antimalarial activity prediction (*−*3.15%) but causes a large drop in predicted step-bunching probability (*−*80.4%), showing that aromaticity can be detrimental if it disrupts the geometry or protonation state of key binding sites.^80,81^

For amines, the pyrrolidines stood out as the dominant determinants of step-bunching. Removing a single pyrrolidine plus its ethyl linker reduced step-bunching by *−*69.2% (Fig. S1b), nearly as much as removing both (*−*84.0%; Fig. 4b), indicating that each pyrrolidine is independently critical for the surface-driven mechanism. Although with a smaller magnitude, the persistent decrease in step-bunching observed when replacing the pyrrolidine nitrogens with *CH* groups (Figs. 4a and S1a) further supports the conclusion that the protonatable nitrogens within these rings are essential for step-bunching crystal growth inhibition.

In contrast, for antimalarial activity, single-nitrogen substitution showed almost no effect from removing nitrogen at most positions (Fig. S1), except for the amine linker, whose removal caused the largest drop among all single-nitrogen changes (Fig. 6b). The amine linker could engage in strong electrostatic interactions with heme propionates^82^ or acidic protein residues,^83^ supporting vacuolar retention, while the pyrrolidines and multiple aromatic regions together could stabilize binding through *π*–*π* stacking,^84,85^ hydrophobic contacts^86,87^, and orientation control.^80,88^

Overall, our theoretical analysis suggests that two key fragments are critical – protonated pyrrolidines (for step-bunching) that may act as cationic anchors to charged step sites,^39,89^ and the chlorobenzene ring (for step-bunching and overall antimalarial activity) which may contribute to additional *π*–*π* stacking and hydrophobic packing to increase residence time at the surface.^84,91^ No other crystal growth inhibitors in our dataset contain a *halogenated benzene* motif (see Fig. 2), despite some being halogenated, suggesting that this aromatic–halogen combination may provide a distinctive structural advantage for PY. Regarding the protonated pyrrolidines, the ethyl linkers may set the pyrrolidines at an optimal distance from the aromatic core, enabling efficient adsorption that immobilizes steps and induces cooperative bunching.^93,94^ For whole-parasite blood-stage antimalarial activity, the mechanism seems to be more complex, involving the combined action of all fragments. In both step-bunching and broader antimalarial activity, there are important position-specific effects, and loss or modification of functional groups at key fragments produces far greater disruption than the removal of comparable groups elsewhere.

### B. Minor roles: halogens, ether, hydroxyl, and phenol groups

Previous studies established that halogens enhance lipophilicity, promote hydrophobic packing and membrane penetration, and strengthen *π*–*π* stacking with heme. Ethers can exert some control over the conformational state of the molecule,^65–67^ whereas hydroxyl groups can tune the polarity;^95^ and both can potentially modulate solubility, pharmacokinetics, and positioning for adsorption.^96^ Phenolic *OH* groups differ from aliphatic *OH* in their greater acidity and resonance stabilization, enabling stronger and more directional hydrogen bonding, *π*–*π* stacking enhancement, polarity tuning, and orientation stabilization at the heme surface.^74–76^

Our theoretical analysis suggests that removal of chlorine from chlorobenzene had minimal effect on either step-bunching or antimalarial activity (see Fig. 6d), indicating that the halogen alone probably plays a supporting role and that its functional contribution is only realized when coupled with the aromatic ring. The aryl portion of chlorobenzene may provide the planar surface for *π*–*π* stacking,^97^ while the chlorine likely enhances polarizability and dispersion interactions, subtly tuning binding strength.^98^ Ether and hydroxyl groups also had minimal impact on step-bunching or antimalarial activity when removed (keeping *>* 88.0–90.0% of PY values), in line with their hypothesized roles as subtle modulators. In the sequential pyrrolidine-removal series, removing phenol after pyrrolidines caused no further meaningful decrease in step-bunching (*−*84.0% vs *−*83.2%; see Fig. 4c), and removing only the phenolic *OH* yielded a similar result (*−*85.4%; data not shown). For antimalarial activity, phenol removal caused a modest decrease (*−*32.9%) compared to removal of the pyrrolidines (*−*23.7%), with most of the effect in the phenol reduction attributable to the *OH* (*−*26.9%). This suggests that the hydroxyl group, similar to the protonatable nitrogens in the pyrrolidines or the chlorine in the chlorobenzene ring, acts synergistically with its local ring environment rather than functioning independently. Both aromatic (chlorobenzene, phenol) and saturated (pyrrolidines) ring frameworks seem to be essential for positioning and stabilizing their respective functional groups – chlorine, hydroxyl, and protonatable nitrogens – in geometries conducive to step-bunching and broader antimalarial activity.

To explain these results, we suggest that the chlorobenzene aromatic–halogen motif probably functions as an integrated unit: the aromatic ring could align with heme for *π*–*π* stacking,^97^ while the chlorine likely enhances polarizability and hydrophobic packing,^98^ together increasing binding strength to the crystal surface.^99–102^ The halogen alone probably lacks this effect,^103,104^ explaining the negligible impact of chlorine removal. Ether and hydroxyl groups, including phenolic *OH*, likely finetune PY’s polarity, solubility, and hydrogen-bonding profile^95,96^ to optimize positioning and persistence at the heme surface or in the digestive vacuole, but they do not appear to drive activity independently. The phenolic *OH* directional hydrogen-bonding potential might help to stabilize the orientation established by primary binding groups,^105,106^ but only when those groups – pyrrolidines and chlorobenzene – are already engaged.

### SUMMARY, CONCLUSIONS AND FUTURE DIRECTIONS

A novel theoretical method of quantitatively analyzing local correlations between the chemical structures and molecular functions is presented. It utilizes a fragment-deconstruction computational analysis that is specifically applied to pyronaridine (PY), a rare small-molecule which uses the step-bunching mode of hematin crystal-growth inhibition as its main tool in fighting malaria. Our method integrates parallel computational metrics for step-bunching probability and MAIP-predicted blood-stage antimalarial activity. By systematically removing, substituting, and modifying functional groups, we distinguished structural motifs with primary, position-specific roles in step bunching and antimalarial activity from those acting mainly in supportive or context-dependent capacities.

Our findings reveal that step bunching activity is highly localized in a small number of structural features, most notably in the pyrrolidines, with the chlorobenzene ring serving as a strong secondary contributor. These fragments appear to act as direct, surface-binding elements, with pyrrolidines functioning as cationic anchors to charged step sites on crystal surfaces and chlorobenzene providing *π*–*π* stacking and hydrophobic packing to increase the residence times. In contrast, antimalarial activity seems to be more distributed, relying on a coordinated interplay between mechanism-driving motifs (e.g., pyrrolidines, chlorobenzene, aromatic–heteroatom scaffolds) and auxiliary pharmacophoric elements (e.g., ether, phenol, halogen) for parasite penetration, digestive vacuole retention, and potential multi-target engagement. The amine linker emerged as the single most impactful protonatable nitrogen for broader antimalarial activity, underscoring the importance of position-specific electrostatic contacts in whole-parasite activity.

The roles of ether and hydroxyl groups—including phenolic OH—appear to be primarily modulatory, finetuning polarity, solubility, and hydrogen-bonding patterns to optimize positioning and persistence at the heme surface or within the digestive vacuole. Although their removal had minimal direct effect on step bunching or antimalarial activity predictions, future studies could probe these groups under conditions where polarity balance or hydrogen bonding is expected to become limiting, potentially revealing context-dependent contributions.

From a design perspective, retaining the step-bunching “core” – pyrrolidines positioned by ethyl linkers and an aromatic–halogen motif – should be a primary goal for preserving irreversible, surface-driven crystal growth inhibition. Peripheral groups can then be strategically tuned with the targeted addition of modulators such as ethers and hydroxyl to improve pharmacokinetics, solubility, or combination therapy compatibility without compromising the geometric and electrostatic features essential for step bunching. This principle may extend beyond malaria to other crystallization-driven diseases, where controlling surface adsorption and defect propagation is therapeutically valuable.

While our theoretical method can quantify the local correlations between structural and functional properties of molecules, it is important to discuss its limitations. The step-bunching probabilities and antimalarial prediction scores are computational proxies, not direct experimental measurements, and may be influenced by model biases, descriptor correlations, or uneven sampling in the training data. Our probability-prediction method, developed from a small dataset of crystal growth inhibitors with well-characterized mechanisms, is an initial effort and requires formal validation against independent experimental benchmarks. Additionally, the present deconstruction approach explores one systematic order of fragment removal, but many alternative deconstruction paths are possible. Specifically, fragments were removed or substituted following a single, predefined sequence in each analog series. We did not exhaustively test all possible orders of removal, so the observed effects may depend on the chosen sequence, especially where fragments exhibit cooperative or non-additive interactions. Fragment “splicing” strategies, such as connecting pyrrolidines directly to the amine linker, substituting nitrogen with methyl groups, or embedding nitrogen into alternate ring systems, but in chemically plausible ways, could reveal further positional and cooperative effects.

Furthermore, the fragmentation and deconstruction approach focuses on discrete structural modifications, which may not fully capture conformational dynamics, solvation effects, or induced-fit changes upon hematin binding; such factors could influence real-world step-bunching behavior in ways not reflected in the static model outputs. The current study examines PY in isolation, yet the mechanistic contributions of individual sub-structures may differ in analogs with altered substitution patterns or in combination therapies where intermolecular interactions and competition for binding sites occur. Moreover, the antimalarial prediction scores from MAIP likely integrate multiple determinants of blood-stage activity—including cellular uptake, metabolic stability, and possible engagement with secondary targets, yet the relative contributions of these processes remain to be experimentally determined.

Future work could integrate molecular dynamics simulations of PY–hematin interfaces under acidic conditions to directly observe binding poses and step-bunching-compatible adsorption patterns. Additionally, experimentally synthesizing targeted PY analogs with selective modification of the pyrrolidine nitrogens, chlorobenzene, or the amine linker could enable more precise causal structure–function analysis across both crystal growth inhibition and whole-parasite assays, refining mechanistic models and guiding rational design of step-bunching-optimized antimalarials with enhanced pharmacological profiles. AFM or in situ microscopy could then confirm whether the predicted high-impact structural features indeed promote step aggregation in live crystal growth environments.

Finally, the modularity of our method makes it generalizable well beyond the present case study. By pairing any mechanistic predictor, whether for crystal growth inhibition, protein–ligand binding, catalytic efficiency, or polymer assembly, with whole-system or application-relevant performance models, fragment-level perturbations can be used to map the structural requirements for mechanism specificity versus overall functional output. In principle, this approach can leverage any predictive model with a well-sampled chemical or material space, the coverage of which can be assessed using similarity metrics against validated reference compounds or structures reported, even without direct access to the training data^107,108^.

Because our theoretical approach is not tied to a particular outcome, it could be deployed across domains ranging from drug discovery and agrochemical optimization to materials science and supramolecular design. Future iterations could integrate automated analog generation, high-throughput multi-endpoint prediction (e.g., stability, solubility, permeability, reactivity, toxicity), and multi-objective optimization algorithms to identify structures that balance competing performance criteria. Such an integrated, mechanistically interpretable framework would enable the rational design of molecules and materials that are simultaneously optimized for mechanistic precision and various applications.

## DATA AVAILABILITY

The data and analysis scripts that support the findings of this study are available from the corresponding author upon reasonable request.

## SUPPLEMENTARY MATERIAL

Additional Analog Analyses: supporting results; Figs. S1-S3); Detailed Methods: Probability Prediction Method, Dataset, Molecule Preparation and Descriptor Calculation, Descriptor Preprocessing and Feature Scaling, Mechanism-Aware Feature Selection via Inter-Class Pairwise Differences, Mechanism Probability Prediction via Distance-Based Scoring, Comparison of Inverse Distance and Softmax Scoring, Interpretation of Model Outputs, Antimalarial Activity (description of the significance of blood-stage antimalarial activity in the context of parasite life cycle and the significance of *Plasmodium falciparum* and how it differs from other *Plasmodium* species)

## ACKNOWLEDGMENTS

The work was supported by the Welch Foundation (C-1559), the NIH (R01GM148537) and the Center for Theoretical Biological Physics sponsored by the NSF (PHY-2019745).

