## Supplementary material for "Physical–Chemical Approach to Identify Local Structural Determinants of Molecular Mechanisms: Case Study of Antimalarial Drug Pyronaridine and Crystal-Growth Inhibition": SI

In this supporting information we present additional methods, figures, tables, and data that were referenced in the main text.

### Additional Analog Analyses

Fig. S1 shows the predicted values for analogs of PY in which individual nitrogen atoms were substituted with $CH$ groups, targeting: (a) a pyrrolidine nitrogen, (c) a pyridine nitrogen attached to an ether, and (d) a pyridine nitrogen attached to an amine linker. Panel (b) represents removal of a single pyrrolidine together with its ethyl linker.

Predicted antimalarial activity was largely not reduced across these analogs, except for analog (d), which was reduced by 20.3% compared to PY. In contrast, the largest decreases in predicted step-bunching probability were observed for substitution of a pyrrolidine nitrogen $N$ with $CH$ (–40.4%) and for removal of the pyrrolidine with its ethyl linker (–69.2%).

Notably, unlike in Figs. 4 and 5, where fragments were removed sequentially to generate analogs of progressively smaller molecular size, the analogs shown here (a, c, d) are of comparable size. As a result, the contributions of nitrogen atoms at different positions can be evaluated more directly, without confounding effects arising from variations in overall molecular size or from the extent of fragment removal in generating each analog.

Fig. S2 shows the predicted step-bunching probability and antimalarial activity for PY analogs generated by sequential removal of major fragments. Removing both a pyrrolidine and its ether linker and a chlorobenzene group (panel a) reduced the predicted step-bunching probability by 69.5% and antimalarial activity by 44.7%. Removing *both* pyrrolidines with their ether linkers and the chlorobenzene group (panel b) produced the largest step-bunching reduction in the series (−86.5%) and a substantial reduction in predicted antimalarial activity (−72.3%).

Further removal of the amine linker (panel c) through substitution of $NH$ with $CH_2$ yielded similar step-bunching reduction (−88.0%) and only a modest additional drop in antimalarial activity (−79.2%) compared to panel b. Removing the phenol group in addition to all previous fragments (panel d) slightly decreased step-bunching probability (−85.9%) and further reduced antimalarial activity (−87.0%) compared to panel c. Panel (e) shows each analog's predicted values as a percentage of PY; the analog represented in (dH) (not

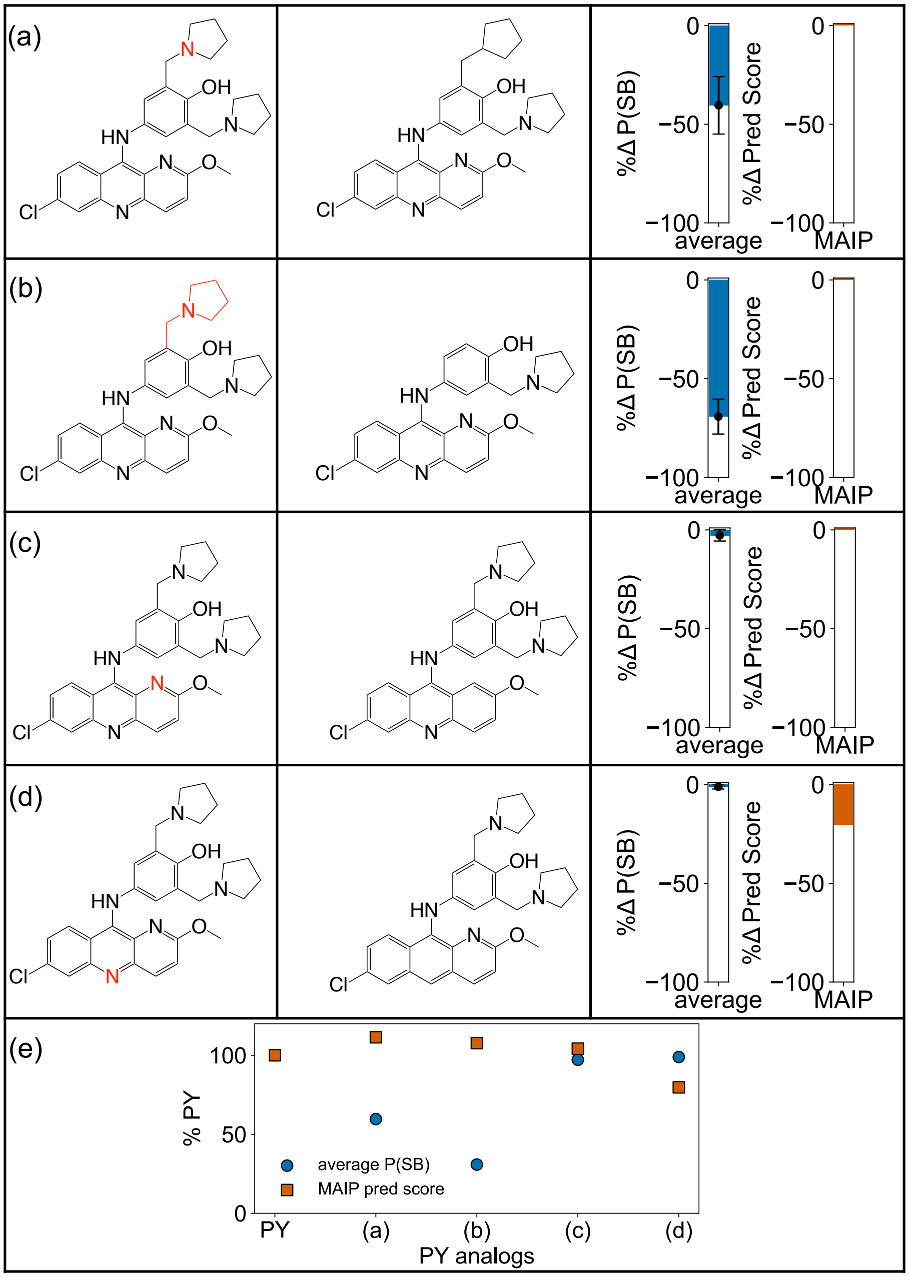


Figure S1: Predicted step-bunching probability and antimalarial activity for pyronaridine (PY) analogs with individual nitrogen ($N$) substitutions or nitrogen-containing fragment removals. (a) Substitution of a pyrrolidine nitrogen $N$ with $CH$; (b) removal of one pyrrolidine and its ethyl linker; (c) substitution of a pyridine nitrogen $N$ (on the pyridine attached to an ether group) with $CH$; (d) substitution of a pyridine nitrogen $N$ (on the pyridine attached to an amine linker) with $CH$; (e) predicted values expressed as a percentage of PY. The left vertical column shows in red the chemical entities removed for creating PY analogs. The middle vertical column shows the considered analogs. Error bars represent range-based variation across two different step-bunching probability prediction methods.

shown in structural form) is derived from panel (d) but with all heteroatoms substituted — $N$ replaced by $CH$ and $O$ replaced by $CH_2$ — which caused almost no change in either predicted value.

As shown in Fig. S2(e), mean predicted step-bunching probability and antimalarial activity both showed sharp decreases from the parent structure (PY) to panel (a). Between panel (a) and panel (b), there was a smaller but still substantial drop — from 30.5% (panel a) to 13.5% (panel b) of PY for step-bunching probability, and from 55.3% (panel a) to 27.7% (panel b) of PY for antimalarial activity. From panel (b) to panel (e), changes were minimal and nonlinear for step-bunching probability (13.5% to 14.7%), whereas antimalarial activity showed a modest but steady linear decline (27.7% to 4.16% of PY).

Fig. S3 presents the predicted step-bunching probability and antimalarial activity for PY analogs with specific functional group substitutions or structural modifications.

Removal of the hydroxyl ($OH$), substitution of the $NH$ in the amine linker with $CH_2$, substitution of all nitrogens $N$ in six-membered aromatic rings with $CH$, and substitution of the $O$ in ether with $\mathrm{CH_2}$ (panel a), resulted in a small decrease in predicted step-bunching probability (−2.38%, 97.62% of PY) but a substantial reduction in predicted antimalarial activity (−44.2%, 55.8% of PY).

Removing the double bonds from every aromatic six-membered ring except the chlorobenzene (panel b) resulted in a small reduction in step-bunching probability (−5.76%, 94.2% of PY), similar to panel (a), but produced a smaller decrease in predicted antimalarial activity (−30.8%, 69.2% of PY) than in panel (a). This suggests that while the aromaticity of non-chlorobenzene six-membered rings may contribute to antimalarial activity, it is less critical than the interactions involving functional groups and heteroatoms perturbed in panel (a).

Replacing both pyrrolidines with pyridines (panel c) caused a dramatic reduction in predicted step-bunching probability (−80.4%, 19.57% of PY) but only a minor change in predicted antimalarial activity (−3.15%, 96.85% of PY), highlighting the pyrrolidines' strong association with step-bunching and minimal contribution to antimalarial activity.

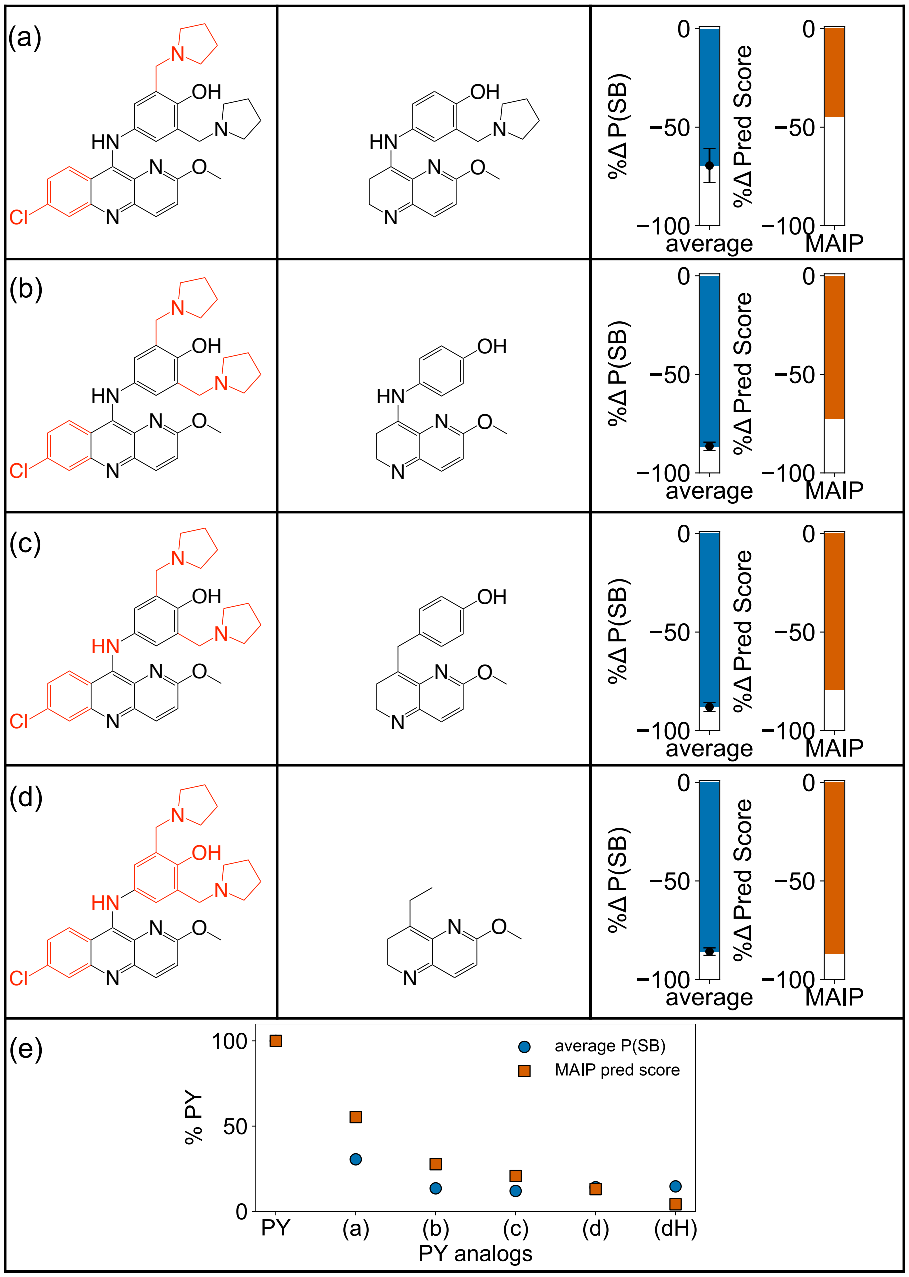


Figure S2: Predicted step-bunching probability and antimalarial activity for pyronaridine (PY) analogs with sequential fragment removal. (a) One pyrrolidine with its ether linker removed and chlorobenzene removed; (b) both pyrrolidines with ether linkers removed and chlorobenzene removed; (c) Analog in (b) plus amine linker removed through substitution of $NH$ with $CH_2$; (d) Analog in (c) plus phenol removed; (e) predicted values expressed as a percentage of PY. Analog (dH) in panel (e) is the analog in panel (d) but with all heteroatoms substituted (N → CH, O → $CH_2$). The left vertical column shows in red the chemical entities removed for creating PY analogs. The middle vertical column shows the considered analogs. Error bars represent range-based variation across two different step-bunching probability prediction methods..

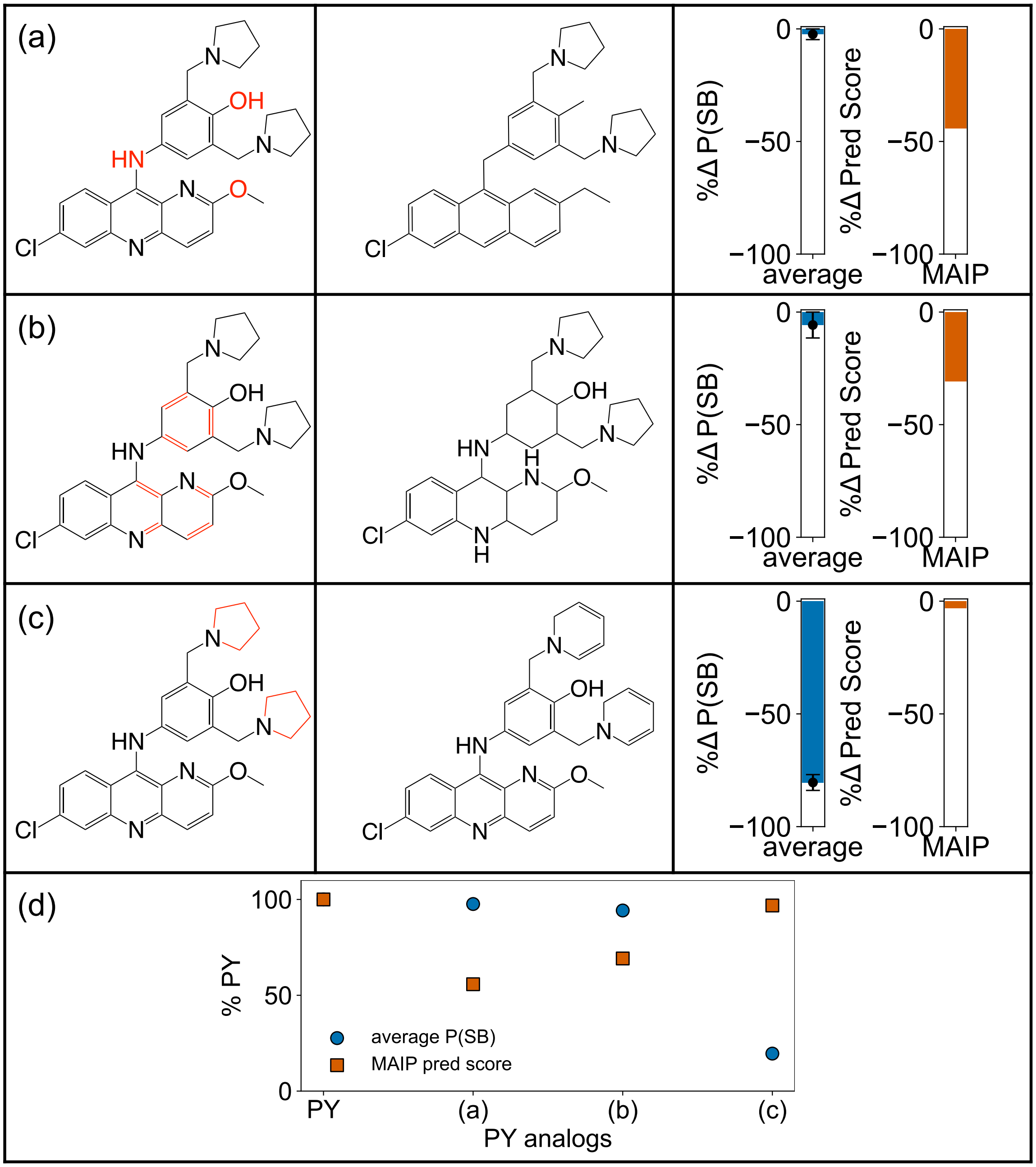


Figure S3: Predicted step-bunching probability and antimalarial activity for pyronaridine (PY) analogs with specific functional group substitutions or structural changes. (a) Removal of $OH$, substitution of the $NH$ with $CH_2$, substitution of nitrogens $N$ in six-membered aromatic rings with $CH$, and substitution of the $O$ in ether with $CH$; (b) substitution of double bonds in all aromatic six-membered rings except chlorobenzene with single bonds to convert aromatic rings to saturated rings; (c) replacement of both pyrrolidines with pyridines; (d) predicted values expressed as a percentage of PY. The left vertical column shows in red the chemical entities removed for creating PY analogs. The middle vertical column shows the considered analogs. Error bars represent range-based variation across two different step-bunching probability prediction methods.

### Detailed Methods

#### Probability Prediction Method

##### Dataset

The dataset consisted of experimentally verified crystal-growth inhibitors of $\beta$-hematin (synthetic hemozoin), each assigned to a specific mechanism class based on atomic force microscopy (AFM) growth assays conducted under physiologically-relevant conditions[1] in prior published studies:[2,3]

- Step-pinners: Quinine (QN), chloroquine (CQ)
- Step-buncher: Pyronaridine (PY)
- Kink-blockers: Amodiaquine (AQ), mefloquine (MQ)
- Passive bystander: Artemisinin (ART)

These assignments reflect direct experimental observation of drug–crystal surface interactions and resulting changes in step advancement behavior. The dataset included only compounds with unambiguous mechanism classification from high-resolution AFM measurements to ensure clean ground-truth labels for model development and interpretation.

##### Molecule Preparation and Descriptor Calculation

Molecular structures and SMILES strings for PY and its analogs were curated from experimental studies on crystal growth inhibition mechanisms. All molecules were processed using RDKit for SMILES parsing, molecular graph validation, and chemical reasonableness checks. The complete set of 2D Mordred descriptors was computed for each molecule. Descriptors with missing values for any molecule were excluded, and molecules for which descriptor calculation failed were removed from further analysis. This process yielded a descriptor matrix of 1425 numeric features.

#### Descriptor Preprocessing and Feature Scaling

To ensure comparability across features and avoid bias in distance-based calculations, all descriptors were scaled to the [0, 1] range using min–max normalization:

$$x' = \frac{x - x_{\min}}{x_{\max} - x_{\min}}. \tag{S1}$$

Scaling was performed with the `MinMaxScaler` from `scikit-learn`, fitted on the cleaned training set and applied to both training and test molecules using identical parameters.

#### Mechanism-Aware Feature Selection via Inter-Class Pairwise Differences

A custom feature selection approach was developed to identify descriptors most discriminative for each of the four crystal-growth inhibition mechanism classes. For a given class $C$, every molecule in the class ($m_i \in C$) was compared to all molecules outside the class ($m_j \notin C$) by calculating absolute differences for each descriptor. For each feature, the smallest of these inter-class differences observed across molecules was recorded, representing the smallest separation from any out-of-class molecule (minimum inter-class distance). Large minimum inter-class distances therefore indicate that a feature provides stronger separation between classes and has greater discriminatory power for a given mechanism. This process was repeated for all molecules ($m_i \in C$) and features. Descriptors were ranked according to the minimum inter-class distances. The top $k$ (typically $k = 5$) consistently identified across members of a class were retained as the class-specific discriminative set, denoted $F_C$.

#### Mechanism Probability Prediction via Distance-Based Scoring

For each mechanism class $C$, the class centroid $\mu_C$ was computed in its class-specific feature subspace $F_C$.

For each feature $f \in F_C$, the centroid coordinate represents the mean of the standardized (values scaled from 0 to 1) descriptor values across all molecules in the in the class:

$$\mu_{C,f} = \frac{1}{|C|} \sum i \in C x_{i,f}. \tag{S2}$$

Given a test molecule $m$, its scaled descriptor vector was restricted to the same feature set $F_C$, forming the subvector $\{x_{m,f} : f \in F_C\}$. The similarity of $m$ to class $C$ was then quantified by its Euclidean distance to the class centroid:

$$d_C(m) = \sqrt{\sum_{f \in F_C} \left(x_{m,f} - \mu_{C,f}\right)^2}. \tag{S3}$$

Here, the projection into $F_C$ simply denotes selecting the corresponding feature components from the full descriptor vector.

Two schemes were used to convert these distances into probability-like scores:

- **Inverse Distance Normalization:**

$$P_C^{(\mathrm{inv})}(m) = \frac{1}{d_C(m) + \varepsilon} \Bigg/ \sum_{C'} \frac{1}{d_{C'}(m) + \varepsilon}, \tag{S4}$$

  where $\varepsilon = 10^{-8}$ prevents division by zero.

- **Softmax Transformation:**

$$P_C^{(\mathrm{softmax})}(m) = \frac{e^{-\beta d_C(m)}}{\sum_{C'} e^{-\beta d_{C'}(m)}}, \tag{S5}$$

  where $\beta$ controls the sharpness of the distribution (default: $\beta = 1.0$).

These probabilities provide an interpretable measure of the likelihood that a molecule acts via each of the four crystal-growth inhibition mechanisms.

##### Comparison of Inverse Distance and Softmax Scoring

While both inverse distance and softmax transformations convert distances into probability-like scores, their mathematical forms lead to distinct behaviors. Inverse distance normalization ($P_C^{(\text{inv})}$) assigns probabilities proportional to the reciprocal of the distance, meaning that very small distances yield disproportionately large contributions to the numerator. As a result, if a test molecule is very close to a single class centroid in its feature space, the corresponding probability can become artificially inflated toward 1.0, even when distances to other classes are also relatively small. This makes inverse distance scores highly sensitive to small absolute differences and can overstate confidence in borderline cases.

In contrast, the softmax transformation ($P_C^{(\text{softmax})}$) applies an exponential decay to the distances before normalizing, controlled by the temperature parameter $\beta$. This ensures that differences in distances are mapped to probabilities in a smoother, bounded manner, reducing the dominance of any single small distance unless the separation is truly large relative to all others. Consequently, softmax scores tend to be more moderate, providing a more conservative estimate of class probability when class distances are comparable.

##### Interpretation of Model Outputs

The mechanism-probability framework described here differs from traditional supervised classification in that it does not attempt to produce a single, discrete mechanism label for each molecule. Instead, it uses a small, experimentally validated reference set to define class-specific descriptor centroids and then assigns relative probability-like scores based on the molecule's proximity to each centroid in a reduced descriptor subspace. This is more akin to a nearest-centroid clustering approach than to a fully trained discriminative classifier.

Given the small size of the mechanism-verified dataset, the absolute probability values should be interpreted qualitatively, as indicators of relative similarity to each mechanism class rather than as calibrated likelihoods. Our primary use of these scores is comparative—to examine how systematic structural modifications shift a molecule's position in mechanism space—rather than to make absolute class assignments.

Because the method is similarity-based and parameter-light, formal cross-validation was not applied; cross-validation on such a small set risks overfitting and would not meaningfully improve confidence in the absolute probability values. Instead, we focus on relative changes in the probability distribution for a given molecule across analog series, which is less sensitive to dataset size and emphasizes interpretable structure–mechanism trends.

#### Antimalarial Activity

Malaria is a parasitic disease caused by protozoa of the genus *Plasmodium* and transmitted through the bites of infected female *Anopheles* mosquitoes. Of the five *Plasmodium* species known to infect humans—*P. falciparum*, *P. vivax*, *P. ovale*, *P. malariae*, and the zoonotic *P. knowlesi*—*P. falciparum* is the most lethal and the primary cause of severe malaria and malaria-related deaths worldwide (see reviews:[4–10]).

The *Plasmodium* parasite undergoes distinct morphological and functional transformations as it alternates between the mosquito vector and the human host. Each form is adapted to a specific role in the life cycle:

1. **Sporozoite stage (mosquito to human transmission):** Sporozoites are elongated, motile forms injected into the human bloodstream during the bite of an infected female *Anopheles* mosquito. Their primary function is to migrate rapidly to the liver, initiating the subsequent stage of infection.

2. **Liver stage (exoerythrocytic stage):** Within hepatocytes, sporozoites transform into rounded, metabolically active forms that undergo repeated asexual replication, producing thousands of daughter parasites termed merozoites. In *P. vivax* and *P. ovale*, a subset of sporozoites becomes dormant hypnozoites, capable of reactivation months to years later and causing relapse; this stage is absent in *P. falciparum.*

3. **Blood stage (erythrocytic stage):** Merozoites are released from the liver into the bloodstream, where they invade erythrocytes. Inside red blood cells, the parasite progresses through three main forms:

- *Ring stage:* Characterized by a ring-like appearance under light microscopy, consisting of a thin rim of cytoplasm encircling a central vacuole.
- *Trophozoite stage:* The actively feeding and growing form, responsible for hemoglobin digestion and production of free heme, which is detoxified via hemozoin crystallization.
- *Schizont stage:* The mature form containing numerous daughter merozoites, which are released upon rupture of the host cell to initiate new cycles of invasion.

The synchronous rupture of infected erythrocytes is associated with the characteristic periodic fever, chills, and anemia of malaria.

4. **Sexual stage (gametocytes):** A subset of intraerythrocytic parasites differentiates into male and female gametocytes, which in *P. falciparum* are crescent-shaped. When ingested by a mosquito during a blood meal, these forms undergo fertilization and sporogony in the mosquito midgut, producing new sporozoites that migrate to the salivary glands, ready to infect another human host.

*P. falciparum* differs from other human-infecting *Plasmodium* species in several key aspects:[9]

- Virulence and mortality: Causes the most severe and often fatal cases of malaria due to high parasite densities, sequestration of infected RBCs in microvasculature, and multi-organ involvement.
- Drug resistance: Has developed resistance to multiple drug classes, including chloroquine, antifolates, and more recently artemisinins, making it a primary target for new antimalarial strategies.
- No dormant liver stage: Unlike *P. vivax* and *P. ovale*, *P. falciparum* does not form hypnozoites, meaning relapse is not a feature—but recrudescence from incomplete clearance can occur.

Pyronaridine (PY) exerts its antimalarial activity during the intraerythrocytic (blood) stages by inhibiting $\beta$-hematin (hemozoin) crystallization, a process essential for detoxifying free heme released during hemoglobin digestion in the parasite's digestive vacuole.

- Life-cycle stage activity: PY has potent activity against trophozoites, the stage with the highest heme turnover, but importantly, also exhibits significant activity against ring stages, including *artemisinin*-resistant Kelch 13 mutant parasites.[10]

- Species coverage: In clinical and in vitro studies, PY is effective against *P. falciparum* and *P. vivax*, including multidrug-resistant strains.[11,12] Its activity against *P. vivax* is particularly valuable in co-endemic regions, where dual-species infections are common and control strategies require broad-spectrum agents.

PY's ability to act at multiple *P. falciparum* life-cycle stages, including ring-stage parasites that are less susceptible to many other quinoline-class drugs, expands its clinical utility, particularly in combination therapies designed to clear resistant infections. Its potency against both *P. falciparum* and *P. vivax* provides a dual benefit in co-endemic settings, reducing the need for species-specific treatments and lowering the risk of treatment failure in mixed infections. Furthermore, targeting the blood stages responsible for pathology ensures rapid clinical improvement, while its long half-life supports extended prophylactic coverage post-treatment.
